# Systematic deconstruction of myeloid cell signaling in tuberculosis granulomas reveals IFN-γ, TGF-β, and time are associated with conserved myeloid diversity

**DOI:** 10.1101/2024.05.24.595747

**Authors:** Joshua M. Peters, Hannah P. Gideon, Travis K. Hughes, Cal Gunnarson, Pauline Maiello, Douaa Mugahid, Sarah K. Nyquist, Joshua D. Bromley, Paul C. Blainey, Beth F. Junecko, Molly L. Nelson, Douglas A. Lauffenburger, Philana Ling Lin, JoAnne L. Flynn, Alex K. Shalek, Sarah M. Fortune, Joshua T. Mattila, Bryan D. Bryson

## Abstract

Myeloid cells are key constituents of tuberculosis (TB) granulomas. They are the major target of pathogen infection and play central roles in pathogen control, antigen presentation, adaptive immune cell recruitment, and tissue homeostasis. However, the role of myeloid cells in TB has been studied largely through *ex vivo* experimental approaches that do not capture the dynamic phenotypic and functional states of these cells in the disease environment. To address this gap, we used a combination of bulk and single-cell RNA sequencing (scRNA-seq), computational modeling, and imaging to define the molecular diversity of myeloid cells in granulomas from *Mycobacterium tuberculosis*-infected nonhuman primates. We observed an increase in myeloid cell diversity in granulomas compared to non-granulomatous lung tissue. This increased transcriptional diversity is defined by a continuum of macrophage differentiation-, metabolism-, and cytokine-regulated transcriptional programs. *In vitro* experimental modeling of monocyte-to-macrophage differentiation in defined cytokine environments implicates differentiation time, IFN-γ, and TGF-β signaling as candidate drivers of macrophage diversity. We next examined the conservation of these populations across additional experimental models of Mtb infection and found myeloid cell subsets enriched across the TB disease spectrum. To further contextualize these responses, we constructed an atlas of myeloid cells across diverse human lung pathologies, finding myeloid cell subpopulations that were similar between TB and other lung pathologies as well as subpopulations that distinguish between diseases. Collectively, this study identifies points of integration between myeloid cell biology in TB granulomas and other lung diseases that can be used for defining the signals that instruct myeloid cell behavior in TB and other diseases, as well as advance myeloid cell-targeted therapies.

## INTRODUCTION

*Mycobacterium tuberculosis* (Mtb) infection is responsible for over 1.5 million deaths and more than 10 million cases of active tuberculosis (TB) annually^1^. Granuloma formation is a pathologic manifestation associated with Mtb infection, and these inflammatory lesions contain a diverse collection of cells, including immune cells of the lymphoid and myeloid lineages^2–7^. Under optimal conditions, the activity of these cells is carefully coordinated, prevents bacterial escape and dissemination, and generates sterilizing immunity that kills Mtb^8^. In less optimal conditions, granulomas serve as local sites of bacterial proliferation and contribute to tissue- and organ-level pathology. Differentiating the factors that lead to these disparate outcomes may lead to improved treatments and vaccines for TB.

Myeloid cells are a cornerstone of the immune response to Mtb and play key roles that span the full course of disease from the initiation of infection and development of pathology to disease resolution^9–16^. The range of myeloid cells found in granulomas is diverse and includes mast cells, eosinophils, neutrophils, and multiple subsets of macrophages^2–4,17–27^. Investigation of the complexity of myeloid cells in granulomas has focused on specific features such as the spatial distribution of subsets or polarization along established axes of inflammation^5,24,25,28–32^. Several of these studies emphasize an important conclusion: specific subsets of myeloid cells can shape the outcome of TB disease^25,30^. Many of these causal relationships, however, have only been defined in the murine model of TB which may not adequately capture aspects of the disease pathology observed in humans.

The extent and genesis of myeloid cell diversity in TB granulomas is unclear. Studies of other lung pathologies or myeloid cell dynamics in other tissues provide models (molecular or conceptual) which may apply to TB granulomas. For example, in lung cancer, tissue resident macrophages support systems-level responses that are distinct from those generated by bone marrow-derived cells^33^; in the bronchoalveolar lavage fluid of individuals with severe cases of COVID-19 disease, alveolar macrophages are depleted, and monocyte-derived macrophages take their place^34^. New conceptual models propose that the differentiation of monocytes recruited to sites of inflammation is shaped by both tissue- and inflammation-derived cues. We hypothesize that the genesis of myeloid diversity in TB granulomas results from the combined activity of myeloid cell recruitment and reprogramming of homeostatic circuits in resident cells in response to infection-associated signals. An improved understanding of granuloma myeloid cell diversity and the molecular circuits that control the emergence of specific myeloid cell states may identify pathways that can be targeted to promote anti-mycobacterial and pro-resolution functions.

Here, we sought to define how Mtb infection reprograms the local myeloid cell landscape. We took advantage of the human-like pathology seen in Mtb-infected cynomolgus macaques – an animal model that develops phenotypically diverse granulomas – to identify how alteration of cellular circuits in granulomas shape myeloid cell identity^2,8,35,36^. Our analysis reveals that macrophages harboring a transcriptional signature of monocyte-derived cells are the dominant constituents of granulomas compared to non-diseased lung tissue which harbor mostly macrophages expressing a signature of tissue-derived alveolar macrophages. Furthermore, our ligand-receptor signaling network analysis indicates that TGF-β and IFN-γ signaling are the major axes of variation in granuloma myeloid cells. We also found similar myeloid cell subsets in different lung diseases but variations in their relative abundance, suggesting the presence of a disease-specific local signals that tune myeloid subset emergence or maintenance. Together, these data highlight the unappreciated phenotypic and functional diversity of myeloid cells in TB granulomas and have implications for developing approaches to control Mtb infection and repair damaged lung tissue.

## RESULTS

### TB granulomas harbor diverse myeloid subpopulations

To examine myeloid cell diversity in TB granulomas, we generated single-cell transcriptional profiles from granuloma tissue from Mtb-infected NHP. Four cynomolgus macaques were infected with a low dose of *Mtb* Erdman (<10 CFU) and necropsied at 10 weeks post-infection for profiling (Fig. 1A, Materials and Methods). Granulomas for analysis were selected based on early detection (∼4 weeks post Mtb challenge) on PET-CT scans. 39 lung granulomas and 4 areas of non-granulomatous lung tissue (1 from each macaque) were sampled. Each sample was analyzed for bacterial burden and gene expression using single-cell mRNA sequencing (scRNA-seq)^37^. Granuloma bacterial burden spanned from 25 to 18,600 CFUs. After applying quality control filters (Methods), 31,198 cellular transcriptomes were generated.

**Fig. 1.**
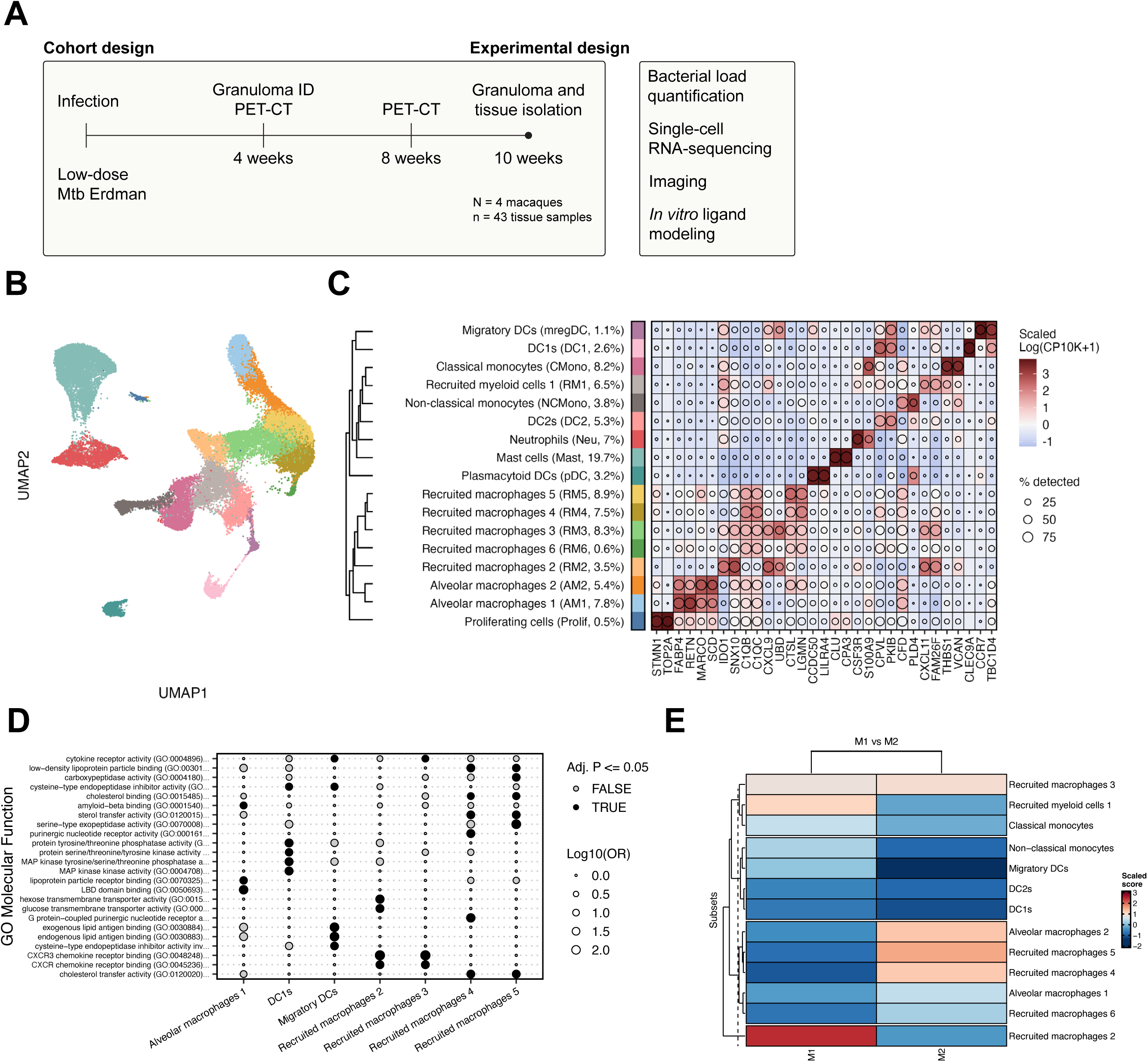
Increased myeloid cell diversity in granulomas. **(A)** Diagrammatic overview of study workflow **(B)** UMAP embedding of integrated cells colored by annotated cell state. **(C)** Clustered heatmap of gene expression, scaled log10(TP10K+1), across cell states. The top 2 markers by AUROC are shown for each state. **(D)** GO molecular function enrichment for the top 50 markers across cell states. **(E)** Hierarchically clustered scaled scores of M1 and M2 transcriptional signatures across macrophages states.

We next integrated these transcriptomes with data from our previously published study of NHP granulomas at 4- and 10-weeks post infection (Materials and Methods, Table S1, Fig. S1-2)^3^. The integrated dataset is composed of 10 macaques, 43 tissue samples, and 41,559 cells (Fig. 1A). Clustering and differential expression analysis of the myeloid cells identified 17 distinct clusters.

Using curated gene signatures for myeloid cells, we identified dendritic cells, mast cells, monocytes, neutrophils, and macrophages (Fig. 1B-C, Table S2-3, Materials and Methods)^3,38,39^. Four dendritic cell populations, including DC1s, DC2s, LAMP3+ DCs, and plasmacytoid DCs were identified. DC1s were defined by *CLEC9A* expression whereas DC2s were defined by *GPR183, CD1C, CLEC6A, CLEC4A* expression. Plasmacytoid DCs had high expression of *LILRA4* and *CCDC50*, and the LAMP3+ DC population resembled previously identified anti-tumor populations based on *CCR7* and *LAMP3* expression^40^. Mast cells were defined by *CLU* and *CPA3* expression. Classical and non-classical monocytes were defined based on *VCAN* and *FCGR3A* expression, respectively. One myeloid population clustered with the monocyte populations but displayed increased expression of other dendritic cell and macrophage genes, such as *IDO1*, *FAM26F,* and *CPVL* and reduced expression of *VCAN* and other monocyte markers. We therefore annotated these cells as recruited myeloid cells (RM1) to highlight the mixed markers^41–43^. Neutrophils were defined by *CSF3R* and *S100A9* expression. Seven macrophage populations defined by *FABP4, MRC1*, *C1QB and CSF1R* expression were identified. Macrophage populations were differentiated by the level of expression of antimicrobial genes, metabolic genes, and metallothionein genes including *CTSB*, *IDO1*, *SOD2*, and *LGMN*. To examine features of macrophage ontogeny, we utilized published signatures of monocyte-derived and alveolar macrophages, scored all macrophage populations according to these signatures, and assigned macrophage class based on signature score^33^.

Gene ontology enrichment using the GO Molecular Function database (Table S4, Materials and Methods) indicated that a range of processes were enriched uniquely in each macrophage subset (Fig. 1D). For example, macrophage populations 4 and 5 had high levels of expression of genes involved in cholesterol metabolism, a key nutrient source for Mtb, while macrophage populations 2 and 3 expressed high levels of genes associated with chemokine receptor signaling^44^. We also sought to contextualize these myeloid cells according to their inflammatory state using published signatures of macrophages stimulated with IFN-γ+LPS or IL-4, reflective of a classical M1 or M2 state, respectively^45^. RM2 cells scored highly for the M1 signature while RM4, 5, and AM2 scored highly for the M2 signature (Fig. 1E). The other populations did not score highly for either signature suggesting that additional signals shape granuloma myeloid cell identity.

In the murine model of TB, interstitial macrophages and alveolar macrophages differ in their antimicrobial capacity^30^. We therefore examined univariate relationships between myeloid population abundance and granuloma bacterial burden. Using a non-parametric Mann-Whitney test, we did not observe a strong association between specific subpopulations and bacterial burden (Table S5). A generalized linear model revealed associations between LAMP3+ DCs and reduced CFU burden as well as an association between mast cells and higher CFU burden as we observed previously^3^ (Table S5).

### Cell recruitment, activation and differentiation underlies the diversity of myeloid cell states in TB granulomas

An emerging model of myeloid cell states within the tissue, such as the lung, involves the dynamic recruitment, activation, and reprogramming of myeloid cells upon deviation from homeostasis^46–48^. We hypothesized that comparing non-granulomatous lung tissue to granulomas would provide insight into the cellular and molecular signals that shape granuloma myeloid cell identity.

We initially hypothesized that the TB granuloma would be associated with less myeloid diversity compared to non-granulomatous lung tissue given dominant inflammatory signaling. We compared myeloid cell diversity in granulomas versus non-granulomatous tissues. Using the Inverse Simpson Index (ISI, range: 1+, higher diversity = higher ISI), we found that granulomas were in fact more diverse (ISI = 9.64) than non-granulomatous tissue (ISI = 6.09) (Fig 2A). We found significant changes in the frequency of specific myeloid populations (adjusted P < 0.05, Methods). Macrophage populations AM1 and AM2 (odds ratio, μOR = 0.152) and monocytes (odds ratio, μOR = 0.51) were enriched in non-granulomatous tissue whereas all other myeloid subsets were enriched in granulomas, except for proliferating cells. We found that macrophages expressing a signature of monocyte-derived macrophages were enriched in granulomas relative to non-diseased tissue consistent with a model of monocyte-mediated replenishment of macrophages in the granuloma niche (Fig. 2B). Plasmacytoid DCs (pDCs), which also showed a dramatic increase in granulomas in our analysis, have been reportedly to be differentially abundant in lung tissue from macaques with latent or active TB disease as well as uninfected versus Mtb-infected mice^49,50^.

**Fig. 2.**
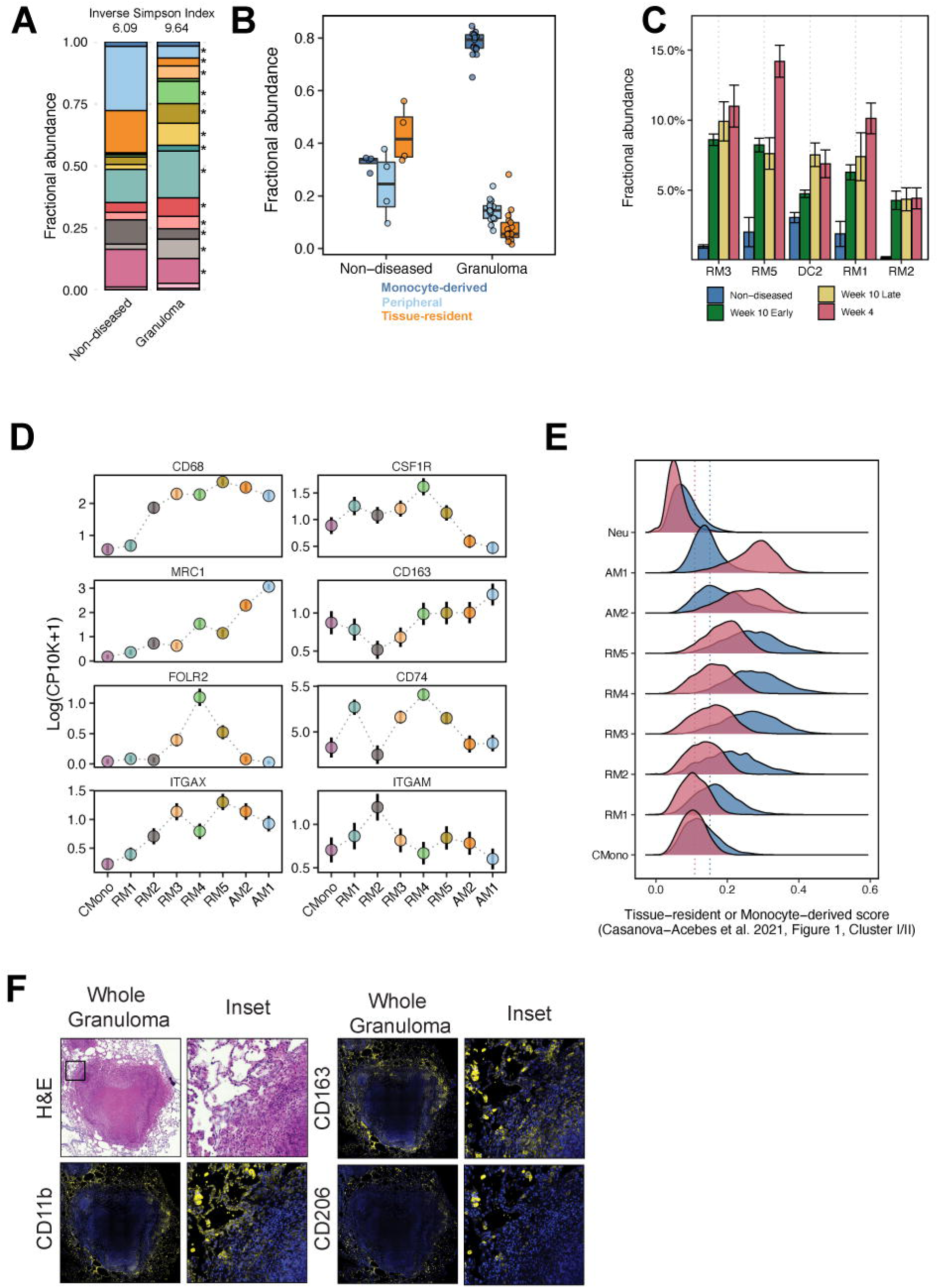
Monocytes and macrophages form a transcriptional continuum aligned with ontogeny. **(A)** Average fractional abundance of cell states between non-diseased and granuloma samples. Inverse Simpson’s Index describes state diversity within each sample type. Asterisk denotes significant change per state in abundance between non-diseased and granuloma samples at adjusted P value < 0.05. **(B)** Fractional abundance of select cell states across week 4, week 10 early, week 10 late, or non-diseased samples. Error bars indicate standard error of the fractional abundance mean. **(C)** Fractional abundance between non-diseased and granuloma samples based on inferred monocyte-derived or tissue-resident ontogeny. **(D)** Log-normalized expression of canonical markers across select monocyte and macrophage markers. Error bar represents standard error of the mean expression per subset. **(E)** Distribution of signature scores across subsets. Signatures are derived from Casanova-Acebes *et al.* 2021 for murine and human monocyte-derived and tissue-resident macrophages. **(F)** Immunofluorescence staining of CD11b, CD163, and CD206 in macaque granulomas.

We next sought to deconstruct other factors that may contribute to granuloma myeloid cell diversity. Our previously published study of NHP granulomas, now integrated here with these new samples, examined how a granuloma’s composition changed at different timepoints post-development so we examined how myeloid diversity changed as a function of granuloma age^3^. Newly-developed granulomas harvested 4 weeks post infection and granulomas that were found at later timepoints by PET-CT imaging and harvested 10 weeks post-infection showed similarly high levels of myeloid cell diversity, whereas non-granulomatous tissue harbored the lowest diversity (Fig. S3). RM1, RM2, RM3, RM5 and DC2 displayed increased relative abundance in week 4 granulomas relative to non-granulomatous tissue, suggesting that these populations may emerge early in the granuloma environment (Fig. 2C).

We then sought to examine monocyte and macrophage diversity using canonical markers traditionally used to define myeloid subsets using flow cytometry (*CD68*, *CSF1R, MRC1* (CD206), *CD163*, *FOLR2*, *CD74*, *ITGAX* (CD11c), and *ITGAM* (CD11b)) (Fig. 2D). We found that *MRC1*, a marker of alveolar macrophages, is increasingly expressed from classical monocytes (CMono) to alveolar macrophages (AM1, AM2), and previous studies have established that monocytes can differentiate into alveolar macrophages^51^. In contrast, *CD68* shows a strong step-like increase in expression from RM1 to RM2, suggesting an inflection point in cell state. On the other hand, *FOLR2*, a marker used for interstitial macrophages and therapeutic target in cancer is expressed highly in RM4 as compared to other populations^52,53^. Consistent with our analysis of individual markers, scoring each of the granuloma monocyte and macrophage populations according to a previously published tissue-resident macrophage signature revealed a gradual increase in expression of the monocyte-derived macrophage score across the recruited macrophage populations (Fig. 2E). These analyses suggest that granuloma myeloid cells include a diversity of macrophage subsets and that recruited monocytes exist on a differentiation spectrum.

We next used immunofluorescence staining to confirm that subset-defining transcripts were translated into proteins by macrophages and to identify where within the granuloma microenvironment these cells are present. We examined expression of CD11b (*ITGAM,* highest in RM2), CD11c (*ITGAX,* highest in RM5), CD68 (highest in RM5), CD163 (highest in AM1), CD206 (highest in AM1), FOLR2 (highest in RM4) and CSF1R (highest in RM4, Fig. 2F, Fig. S4) and found unique expression patterns as well as overlap between these markers. We found that AMs expressing CD206, FOLR2, and CSF1R were abundant in the granuloma-adjacent lung and that few of these cells had infiltrated into the granuloma. In contrast, CD163 and CD11b were expressed by granuloma-adjacent AMs and macrophages in the granuloma’s lymphocyte cuff. CD11c was the most broadly expressed macrophage-associated antigen and was expressed by AMs and macrophages in the histologically-defined epithelioid macrophage region. CD68, which is often used as a general macrophage marker, was most strongly expressed by cells in the epithelioid macrophage region, especially by the cells adjacent to the caseum. Taken together, these protein-level data support our transcriptional analyses by showing the complexity of granuloma-associated macrophage populations in a spatial context.

### *In vitro profiling* reveals dominant variance induced by time, IFN-γ, and TGF-β

Thus far, we verified that diversity in the granuloma could be driven by variations in the abundance of myeloid subtypes, monocyte infiltration, differentiation, and changes in cell state. Recent work on macrophage ontogeny and monocyte-macrophage dynamics has emphasized the influence of environmental cues on myeloid cell phenotype in the tissue^41,43,48,54–56^. Ligand-receptor interaction prediction methods such as NicheNet offer a technique to predict signals potentially responsible for the transcriptional profile of cell populations of interest. To generate testable hypotheses about ligand signals in the granuloma, we used NicheNet focusing on differentially expressed genes within RM2, RM4, and AM1 populations (Table S6, Materials and Methods)^57^. We focused on these populations because they showed the most distinct transcriptional and functional enrichment profiles that were not well-explained by any of the analyses above. NicheNet analysis predicted several cytokines with potential activity in RM2, RM4, and AM1 cells including TNF-α, IL-13, IL-15, IL-1β, TGF-β, IL-6, and IFN-γ, which have been previously detected in TB granulomas (Fig. 3A)^58–60^. TNF transcript was predominantly detected in myeloid cells, whereas TGF-β was detected in NK cells, T cells, DC2s, and proliferating cells (Fig. S5).

**Fig. 3.**
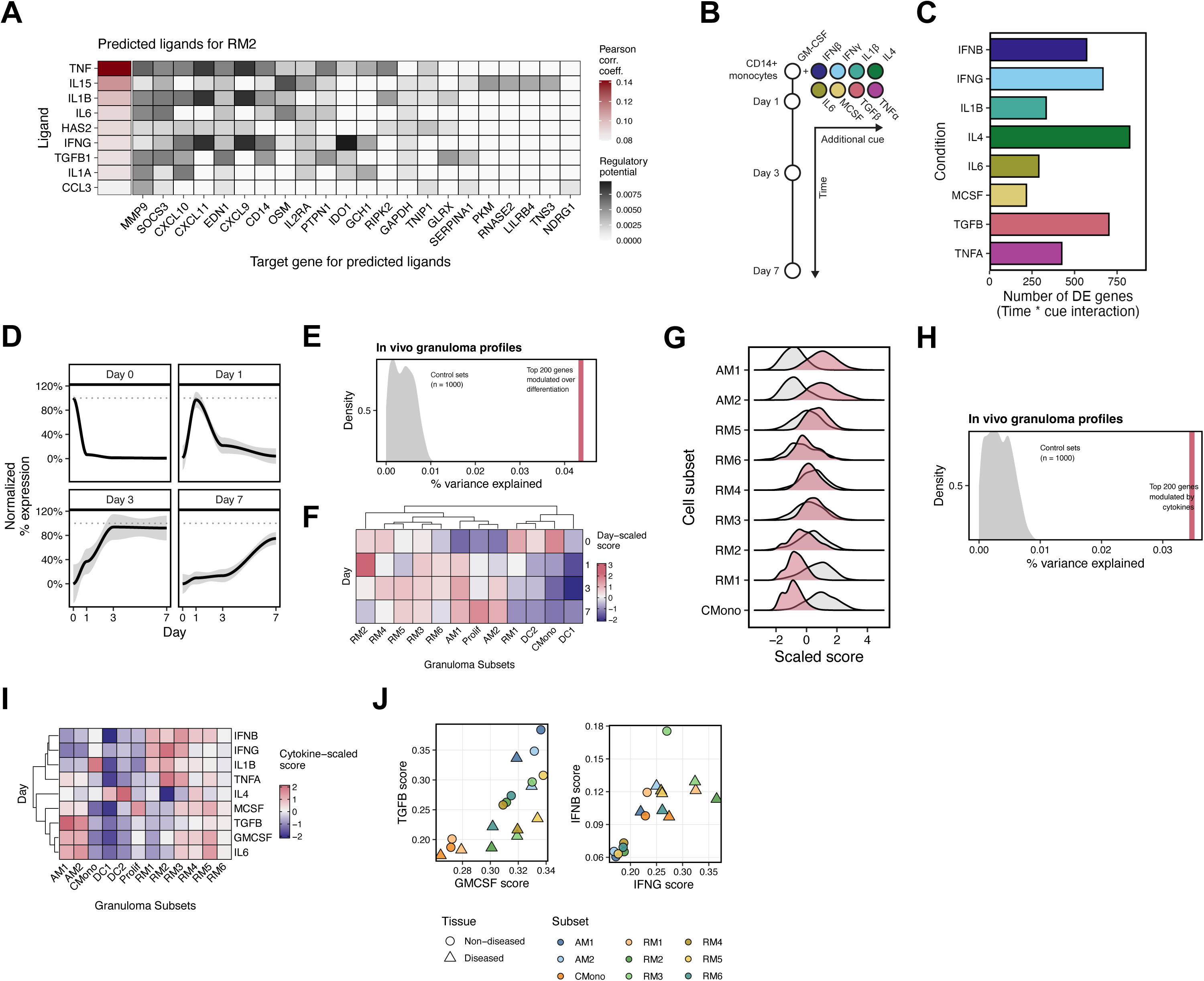
*In vitro* differentiation and stimulation describes *in vivo* transcriptional variability. **(A)** NicheNet predicted ligands based on genes differentially expressed by the RM3 state. **(B)** Overview of experimental setup for *ex vivo* primary macrophage culture, stimulation, and sampling. **(C)** Number of differentially expressed (DE) genes based on time * cue interaction model for each condition. **(D)** Normalized expression of DE genes for each day expressed across the timepoints. **(E)** Percent of variance explained in NHP data by top 200 genes associated with time, based on significance, compared to distribution of 1,000 random gene sets. **(F)** Scaled score for each day signature across NHP myeloid states. **(G)** Distribution of scaled day 1 and day 7 signature scores across NHP myeloid states. **(H)** Percent of variance explained in NHP data by top 200 genes associated with cytokine stimulations, compared to distribution of 1,000 random gene sets. **(I)** Scaled score for each cytokine signature across NHP myeloid states. **(J)** Select cytokine signature scores pseudobulked across subsets between non-diseased and granuloma samples.

Given that the data that inform NicheNet predictions are not derived solely from myeloid cells, we sought to enhance our study of cytokine signals that shape myeloid cell state, by performing time-resolved *in vitro* stimulation experiments of myeloid cells. Given the enrichment of monocyte-derived macrophage signatures in the granuloma, we focused on monocytes and monocyte-derived cells. We and others have previously utilized transcriptional profiling to define the acute response to macrophage stimulation with diverse ligands^61^. To build upon these previous studies and in recognition that monocytes recruited to sites of disease often differentiate in the presence of multiple ligands concurrently, we sought to model the monocyte response to tonic cytokine signals associated with differentiation and granuloma residency. We utilized classical human CD14+ monocytes from peripheral blood as our experimental monocyte source. To examine the contribution of time, we generated samples at multiple time points (0, 1, 3, and 7 days). These studies resulted in the generation of 200 unique RNA-seq samples which we analyzed using Prime-seq, a high-throughput bulk RNA sequencing technique (see Materials and Methods)^62^. To simulate the complexity in lung granulomas, we combined GM-CSF with each of the following ligands: IFN-β, IFN-γ, IL-1β, IL-4, IL-6, M-CSF, TGF-β, and TNF-α (Fig. 3B). Given the necessity and influence of GM-CSF on alveolar macrophage development, ligands were added immediately at day 0 along with GM-CSF, which resulted in robust transcriptional changes over time^63–66^ (Fig. 3C).

We next sought to evaluate how the diverse signals we modeled in our *in vitro* experiments (time and ligand identity) were reflected in the transcriptional signatures observed *in vivo.* We first focused on gene programs that describe the temporal axis of monocyte (day 0) to differentiated monocyte-derived cell (day 7). We defined gene signatures based on differential gene expression at each time point (0, 1, 3, 7 days, Materials and Methods). (Fig. 3D). We identified several time-dependent gene sets: genes that are downregulated following day 0, genes that are induced by day 3 and remain highly expressed at day 7, and genes that are gradually induced over the course of 7 days. Genes in the day 0 signature included *S100A8, CD93, CD14* while genes in the day 7 signature included *MAF, ALOX15B,* and *ITGB5.* We then asked how much variance *in vivo* is explained by the gene sets associated with *in vitro* time course study. We found that the gene sets changing over time *in vitro* explained a significant portion of the variance *in vivo* (Fig. 3E, P < 2.2e-16, Materials and Methods). To identify which time points *in vitro* resemble the *in vivo* subsets, we scored the subsets based on the day-specific gene sets (Fig. 3F). The day 0 gene signature on was most highly expressed by classical monocytes. The day 1 gene signature was expressed most highly by the RM2 subset. The day 7 signature was highly expressed by AM1, AM2, and proliferating cells. Other RM subsets show mixed scoring across day 0 to day 7 consistent with an intermediate phenotype. As an alternative strategy to visualize these trends, we scored granuloma myeloid cells according to the day 0 and day 7 scores and visualized their distributions as a histogram. AM1 and AM2 cells scored higher for the day 7 signature than the day 0 signature. By contrast, classical monocytes (cMono) and RM1 scored higher for the day 0 signature than the day 7 signature (Fig. 3G). Taken together, these data reinforce that granuloma myeloid cells exist on a spectrum of differentiation^41,47^.

We next asked how ligands predicted by our NicheNet analysis and modeled *in vitro* using monocyte-derived cells aligned with variation in myeloid cell gene expression *in vivo.* Like our analysis of temporal signatures, we defined gene sets that describe each ligand using differential expression (Materials and Methods). Like the time-dependent gene sets, the ligand gene sets explain a significant amount of variance *in vivo* relative to random control gene sets (Fig. 3H). We next scored the *in vivo* subsets according to the *in vitro* ligand signatures and compared their relative scores across subsets. GM-CSF and TGF-β gene signatures are most expressed in the AMs with decreasing relative expression to classical monocytes; this is consistent with previous studies demonstrating the requirement of GM-CSF and TGF-β in alveolar macrophage development (Fig. 3I)^63,64,67^. IL-4 signatures associated with the DC subsets^68^. TNF-α and IL-1β signatures showed more distinct subset expression whereas IL-6 and IFN-β showed similar, correlated trends with TGF-β and IFN-γ, respectively (Fig. 3J).

We next visualized sites of *in vivo* TGF-β and IFN-γ signaling by staining granulomas for phosphorylated SMAD3 (pSMAD3) and phosphorylated STAT1 (pSTAT1). We used CD11c as a marker for macrophages based on our prior work showing this marker’s broad expression across subsets (Fig. S6, Fig. 2F, Fig. S4). We found that pSMAD3 signaling was widespread throughout granulomas, including in macrophages (Fig. S6, magenta), whereas cells regulated by STAT1 were less common. Three STAT1 phenotypes were noted in CD11c+ macrophages including pSTAT1-negative cells (phenotype 1), cells with intranuclear pSTAT1 (phenotype 2), and cells with cytoplasmic but not intranuclear pSTAT1 (phenotype 3). Each phenotype could be found in granulomas from macaques at 4- and 10-weeks post infection. pSTAT1-negative macrophages were often found in the granuloma’s lymphocyte cuff region whereas macrophages with intra-nuclear pSTAT1 were present in closer proximity to necrosis. The phenotype 3 macrophages with cytoplasmic pSTAT1 were similar in size and appearance to alveolar macrophages and were present as clusters adjacent to or embedded within the granuloma’s lymphocyte cuff. Taken together, these data suggest that TGF-β and IFN-γ regulated macrophages are distinct subsets of cells that occur in different granuloma regions, potentially with different functional consequences for granuloma-level homeostasis and bacterial control.

### Alignment of myeloid states across species and related pathologies reveals conserved subsets

The abundance of publicly available scRNA-seq data across diverse lung pathologies and other diseases inspired us to examine the possibility that the transcriptional subsets we identified in NHP granulomas were similarly observed across other lung pathologies. To test this possibility, we generated a human lung atlas of myeloid cells (>300,000 cells across 394 samples) in the lung representing diverse pathologies (Materials and Methods, Fig. 4A, Table S7-10)^69^.

**Fig. 4.**
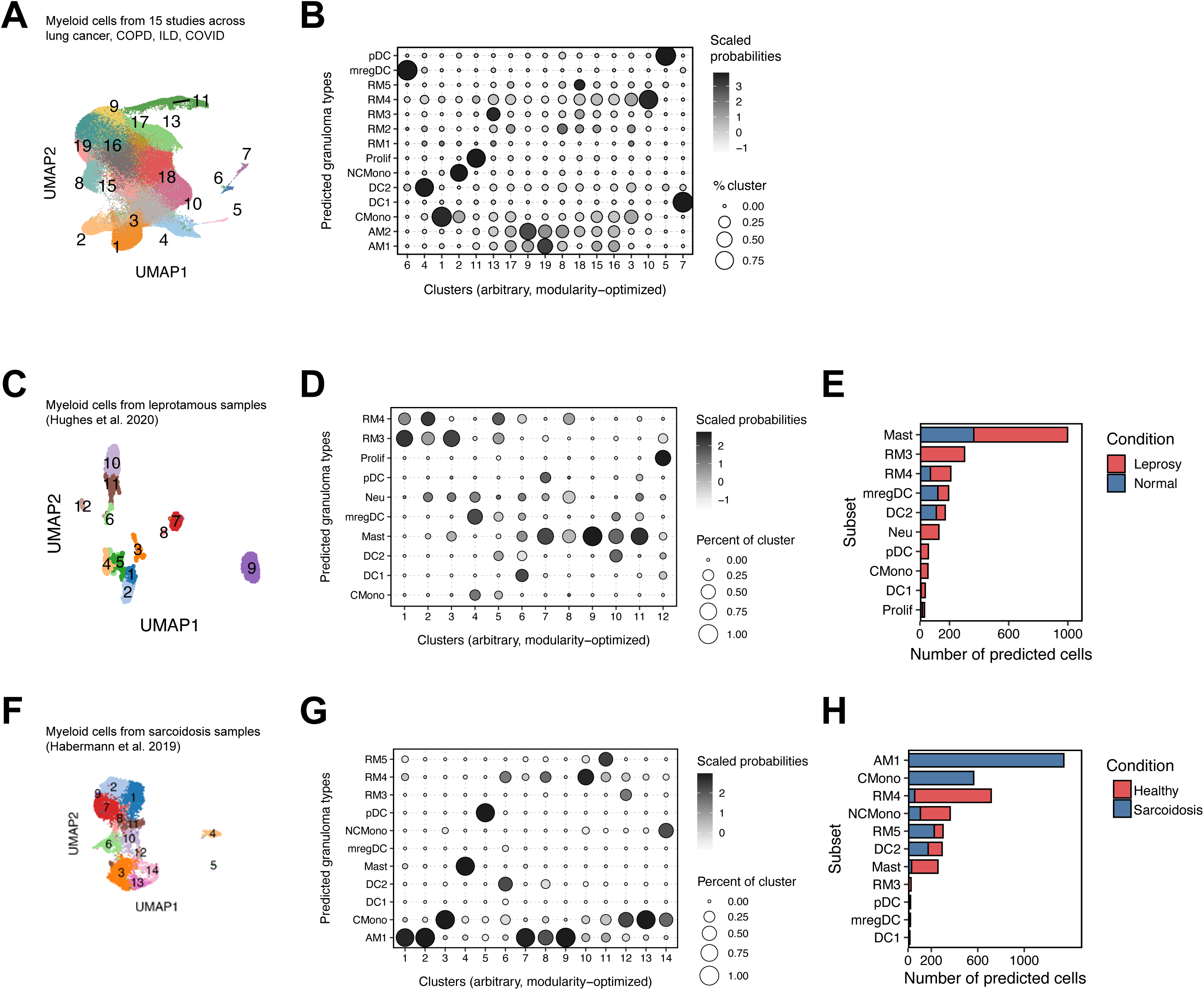
Myeloid human lung atlas suggests conserved disease-induced diversity and transcriptional states. **(A)** UMAP projection of integrated human lung atlas myeloid cells **(B)** CellTypist classification of cell populations across this study and human lung myeloid atlas **(C)** Reanalysis (UMAP) of leprosy granuloma samples from Hughes et al. **(D)** CellTypist classification of cell populations across this study and Hughes et al. **(E)** Comparison of cellular subset abundance in Hughes et al across healthy and granuloma skin samples. **(F)** Reanalysis (UMAP) of sarcoidosis granuloma samples from Habermann et al. **(G)** CellTypist classification of cell populations across this study and Habermann et al. **(H)** Comparison of cellular subset abundance in Habermann et al across healthy and sarcoidosis granuloma samples.

We first asked if there were any unique myeloid cell populations in TB granulomas that were not observed in other pathologies. To compare transcriptional subsets, we utilized Celltypist, a previously established computational framework for scRNA-seq cell type annotation (Materials and Methods)^70^. We trained a CellTypist model on our cynomolgus granuloma data and predicted the granuloma cluster labels within the human lung atlas (Fig. 4B)^38^. We observed strong mapping of most populations, including LAMP3+ DC, DC2, RM3, RM4, RM5, AM1, and AM2 (Fig. 4B). Predictions for RM1 were relatively weaker. Although RM6, defined by metallothionein genes like *MT1X*, failed to generate any predictions, a clear metallothionein-defined cluster (Cluster 17) was identified in the pan-lung pathology atlas. The weak mapping of RM1 may reflect a unique population of granuloma myeloid cells, or it may reflect the dynamic nature of TB granulomas where an immature population of myeloid cells is continually recruited to the granuloma in contrast to the other diseases analyzed.

Our comparison of non-diseased and granuloma tissue revealed a trend involving a reduction in alveolar macrophages and expansion of specific myeloid populations in granuloma tissue. We next sought to test if these trends generalized to the other diseases in our lung atlas. When comparing control versus diseased samples present in the human lung atlas, we saw a significant decrease in the AM1 score and significant increases in LAMP3+ DC, RM3, and RM4 scores in diseased samples overall, consistent with a model where monocyte-derived cells alter the myeloid compartment during disease (Fig. 4C).

Our observation of similar myeloid cell populations between a pan-lung atlas and granulomas next inspired us to investigate specific comparisons between our study and other published studies of mycobacterial disease and granulomatous pathologies.

We compared our cynomolgus macaque study to a previously published study of lung samples from rhesus macaques with latent or active infection^49^. Again, we used Celltypist. We observed robust mapping and subset identification between mast cells, plasmacytoid DCs, and cDC1s across the two datasets (Fig. S7A). We observed more nuanced mapping between macrophage populations. For example, the Alveolar TREM2+ population from the rhesus study mapped to multiple populations in our study (RM4, RM5, and RM6) (Fig. S7A). We next sought to examine the association of specific myeloid subsets from our study with the TB disease status as investigated in the rhesus study. pDCs were elevated in active infection, as described previously, in addition to RM1/RM2/RM3 populations which mapped to their “Alv IFN signature” (Fig. S7B). The alveolar macrophages AM1 and AM2 were more frequent in latent disease consistent with a model where active TB disease alters the lung immune landscape, similar to what is observed in COVID-19^34^.

We next compared the macrophage subsets we identified in NHP with macrophage subsets identified in the lungs of C57BL/6J mice following infection with 1,500 Mtb CFU (Fig. S8A)^71^. Celltypist mapping failed to predict more than one broad cell type, so we modified our analysis to comparing and scoring 1:1 ortholog genes as previously done (Materials and Methods)^72,73^. Non-macrophage subsets and proliferating cells were generally well-aligned (Fig. S8B-C). As previously observed with conservation of macrophage subsets across between mice and humans, concordance between mouse and cynomolgus macrophage subsets were mixed^72^. There was significant similarity between the *Nos2*-expressing IM1 and IM3 populations with RM2 and the *C1QA*-expressing IM2 population with RM4 and RM5.

We next examined the relationship between NHP TB granulomas and other granulomatous diseases. We first compared our myeloid subsets to those observed in leprosy, a skin disease whose causative agent is a different mycobacterial species, *Mycobacterium leprae*^37,74^. Not surprisingly, we observed strong mapping with mast cells and no mapping with alveolar macrophages (Fig. 4D-E). We observed mapping of a limited number of recruited macrophage subsets (RM3 and RM4) to leprosy granulomas. RM3 and RM4 cells were more abundant in leprosy samples than normal skin suggesting that these populations are similarly enriched is a diseased environment (Fig. 4F).

Sarcoidosis is a condition that results in granulomas in the lung and other tissues, and the etiology of sarcoid granulomas is still poorly understood^75^. We asked whether these two granulomatous diseases might have cellular features that distinguish between them^76^. We performed mapping using Celltypist and observed consistent, strong concordance between mast cells, pDCs, and monocytes (Fig. 4G-I). Neutrophils, DC1s, LAMP3+ DCs, RM1 and RM2 largely failed to map to sarcoidosis cells. Notably, RM1 and RM2 cells are marked by high expression of *IDO1*, *CD274* and *CXCL9*. A previous protein-centric study comparing human TB and sarcoidosis granulomas similarly observed an absence of macrophages co-expressing PD-L1 (CD274) and IDO1 consistent with the observations made by imaging mass cytometry^2^. We hypothesize that the absence of neutrophils, DC1s, LAMP3+ DCs, RM1 and RM2 reflects an absence of signals these cells need for recruitment to and differentiation in granulomas. Together, these comparative analyses reveal that granuloma myeloid cells share similarities with other lung pathologies which may facilitate the repurposing of myeloid-targeted therapies in TB as well as mechanistic dissection of the signals that support the generation of these cellular states.

## DISCUSSION

Myeloid cells play a critical role in TB pathogenesis from initiation to resolution^15,25,77^. The function of myeloid cells in granulomas is central to the trajectory of disease. Using a combination of experimental and computational techniques, we defined the transcriptional diversity of myeloid cells in the NHP TB granuloma. We found that granuloma myeloid cells are not a monolith and that cells harboring signatures of monocyte-derived cells are the dominant myeloid cell constituent of granulomas. Many of these myeloid cell populations are detectable as early as 4 weeks. Unlike the mouse model, we did not identify a univariate relationship between cellular subsets and bacterial control. We found that signatures of myeloid cell age and IFN-γ and TGF-β signaling explained a significant component of the *in vivo* transcriptional heterogeneity of granuloma myeloid cells. Lastly, by comparing TB granuloma myeloid cells to other lung pathologies, we found that TB granulomas harbor myeloid cell subsets that are transcriptionally similar to other lung pathologies. Disease-specific comparisons between TB and sarcoidosis granulomas identified cellular features that distinguish between these two types of granulomas.

Myeloid cells integrate diverse signals (ontogeny, soluble cues, time) to shape their identity. Models of macrophage cell states in TB granulomas have historically focused on polarization along an M1-M2 axis, their spatial localization in granulomas, and a small number of canonical markers. In this study, we expand this model to place macrophages on a spectrum from classical monocytes to tissue-resident alveolar macrophages, with each subset being characterized by a distinct transcriptional profile. By comparing to non-diseased lung tissue from the same animals, we demonstrate how the lung tissue niche is remodeled locally in granulomas^33,41,47,78,79^.

*In vitro* profiling further confirmed this spectrum by identifying the mixture of differentiation and cytokine factors that partially describe *in vivo* heterogeneity. Pairing computational predictions of cytokine activities, *in vitro* validation, and imaging of transcription factors that are phosphorylated in response to TGF-β and IFN-γ signaling, we observed that transcriptional variation in the myeloid compartment was associated with variation in IFN-γ and TGF-β signaling among others. The heterogeneity and combination of cytokines measured in granulomas support the idea that granuloma cells entering the microenvironment experience a complex mix of signals^59,60^. Future granuloma myeloid cell phenotyping should incorporate markers beyond canonical macrophage markers and aim to distinguish between monocyte-derived and tissue resident macrophages^80–82^. Based on our findings, candidate markers for expanded protein-centric panels should consider including markers such as NR1H3, CEBPB, CLEC4E, FOLR2, and TREM2 to better define macrophage populations in the granuloma^83–86^.

Our data revealed a macrophage population, RM2, which was high in *IDO1*, *CXCL9*, *CXCL10*, and *CXCL11* expression. RM2 was present across all cohorts and significantly increased in granuloma samples. This population was also high for Mincle (*CLEC4E),* a receptor for mycobacterial ligands^87^. RM1 and RM3 displayed similar, albeit with lower expression of these key features, suggesting that they may be at a different activation or differentiation stage than this population. Metabolically, this population was uniquely high in tryptophan and glycolysis-related pathways and displayed high *STAT1*, *NFKB1*, and *CEPBD* activities. We hypothesize this population represents an immunoregulatory subset composed of recently recruited and immature macrophages. This population shares various features with similar cells described as key mediators of *Salmonella* infection and *Mtb*-infected cells^42,88^. The consequence of this population in the microenvironment remains paradoxical. The expression of tryptophan metabolism and *CEBPD* activity also suggests an immunoregulatory role but the combination of IFN-γ and TNF-α has been noted to drive inflammatory cell death and tissue damage^89–93^. Interrogating the consequence of this population at the site of the granuloma may identify a balance of functional roles this population performs.

Interactions between alveolar macrophages and Mtb are one of the earliest detected interactions between Mtb and the host^94^. While alveolar macrophages are a gateway to the lung early in infection, our data show that monocyte-derived cells are the major contributors to the macrophage compartment in the granuloma. Recent studies in other lung diseases emphasize the importance of monocyte-derived cells. For example, bronchoalveolar lavage samples from individuals with severe COVID-19 disease have a decreased frequency of tissue resident alveolar macrophage and an increased frequency of monocyte-derived cells^34^. It was hypothesized that this increased frequency of inflammatory monocyte-derived cells may be associated with worsened outcomes due by recruiting inflammatory monocytes and neutrophils. The role of monocyte recruitment in TB disease is nuanced and has not been rigorously examined in the NHP model of TB disease. In murine studies, monocyte recruitment has been experimentally explored through the utilization of mice deficient in the chemokine receptor, CCR2^95,96^. The conclusion from these studies is that at high doses of Mtb, loss of CCR2 has a dramatic impact on susceptibility while CCR2 appears dispensable at low doses of Mtb challenge. It is difficult to predict how perturbation of monocyte recruitment in TB disease may impact granuloma formation or TB disease outcome, especially given the diversity of monocyte-derived cells in granulomas. A previous murine study does suggest that monocyte-derived cells have higher antimicrobial potential than alveolar macrophages while a different study suggests that human monocytes have reduced capacity to control Mtb growth compared to monocyte-derived or alveolar macrophages^30,97^. Future experimental studies in NHP may be poised to define a functional role for specific populations of myeloid cells in granuloma function as several myeloid subpopulations were defined by genes encoding cell surface proteins suggesting the potential for antibody-mediated cell depletion. These markers include *CD36*, *CLEC9A, FOLR2, MRC1, MS4A7,* and *SLAMF7* among others.

Recent studies in lung cancer suggest that monocyte-derived cells in the lung may have immunosuppressive functions^33^. TREM2+ monocyte-derived macrophages have become the subject of intense study, in part inspired by a large body of literature on TREM2 in the context of microglia function in the brain. Loss of TREM2 expression or activity has been shown to reduce tumor burden in several models of lung cancer^84,98^. Our data highlight the existence of similar transcriptional populations of TREM2+ cells in TB granulomas. A recent study suggests that TREM2+ macrophages result from the efferocytosis of cellular debris, and it is appealing to consider the contributions of cell death in TB granulomas, which has been widely documented, as a potential driver of this population of cells^98^. While TREM2+ macrophages have been implicated in immunosuppression, it will be necessary to examine whether they play a similarly immunosuppressive role in TB granulomas. More broadly, it will be valuable to explore new experimental perturbations in non-human primates to regulate monocyte-derived cell function, isolate them from granulomas, or model their function with novel *in vitro* models.

Our analyses across other scRNA-seq profiles of lung diseases provides additional experimental support of many of the conclusions made with TB granulomas. Firstly, our observation of similar populations across diseases enhances our confidence in the identification of these transcriptional subsets. The observation of the shift from tissue-resident alveolar macrophages to monocyte-derived macrophages across diseases emphasizes the importance of monocyte differentiation and recruitment as major events that may shape the course of lung diseases. Notably, RM1 and RM2, identified in TB granulomas, weakly mapped to populations in the human myeloid lung atlas. We hypothesize that this may be due to the temporal nature of sampling in these datasets which were generally late-stage fibrotic diseases and cancer; however, future studies should seek to determine if cells resembling the RM1 population is present in other diseases. It has recently been hypothesized that a transitional macrophage population “TransMac” exists during disease. Pseudotime analyses and experimental examination of the myeloid cell populations in the human lung atlas may resolve whether these intermediate populations differentiate into bona fide tissue-resident macrophages or preserve their intermediate state^47,79^.

Recent studies highlight a role for eosinophils in modulating infection and macrophage function^17,18^. In our study, we did not detect the canonical marker defining eosinophils, *EPX*, to any significant degree. Alternative scRNA-seq technologies may better facilitate their capture and analysis^99,100^. Lastly, our *in vitro* studies were limited to a small number of cytokines for experimental feasibility subset of cytokines. The pleiotropic nature of several cytokines included, such as IL-6, further complicate these efforts^101^. Biologically, there may be multiple sets of cytokines that can generate a given cell state and computationally, recovering the stimulation history of cells in tissue is difficult; however, studies with tissue resident alveolar macrophages and cytokine neutralization studies *in vivo* may help disentangle this complexity in the future.

In summary, our integrative and comparative investigation detailed the myeloid cell states in the granuloma microenvironment and across similar pathologies. By identifying and contextualizing these newly identified macrophage populations, we compiled significant evidence for acute, pathology-associated recruited macrophage states (primarily RM2, RM3, and RM4). Better understanding how these cell states modify the adaptive cell compartment will help differentiate the beneficial and pathogenic roles they may play at different points in infection. Taken together, our data substantiate a highly dynamic and microenvironment-driven monocyte-to-macrophage compartment that shares features across diseases and models. This framework has far-reaching implications and suggests the ability to co-opt biology across diseases as our understanding of their dynamics spatiotemporally and ability to therapeutically target these cells increases. Building complete models of macrophage state across perturbations–such as genetic knockouts, cytokines, cellular depletions, and vaccines–will enable rational dissection of the immune responses behind effective vaccines and host-directed therapies in TB.

## SUPPLEMENTARY MATERIALS

### Materials and Methods

Fig. S1. Technical assessment of ambient and batch effects. Fig. S2. Overview of week 10 cohort 2 infection.

Fig. S3. Inverse Simpson’s Index across sample types.

Fig. S4. Immunofluorescence staining of myeloid cell markers in macaque granulomas. Fig. S5. NicheNet analysis of cytokine production.

Fig. S6. Immunofluorescence staining of pSTAT1 and pSMAD3 in macaque granulomas. Fig. S7. Alignment of NHP states with NHP states in Esaulova et al.

Fig. S8. Alignment of NHP states with murine states. Table S1. Macaque and sample metadata.

Table S2. Cellular metadata. Table S3. Cell markers.

Table S4. Cluster gene enrichment. Table S5. CFU associations.

Table S6. NicheNet activities. Table S7. In vitro samples.

Table S8. Public studies used for comparisons. Table S9. Atlas metadata.

Table S10. Atlas markers.

## MATERIALS AND METHODS

### Ethics statement

All experimental manipulations, protocols, and care of the animals were approved by the University of Pittsburgh School of Medicine Institutional Animal Care and Use Committee (IACUC). The protocol assurance number for our IACUC is D16-00118. Our specific protocol approval numbers for this project are 18124275 and IM-18124275-1. The IACUC adheres to national guidelines established in the Animal Welfare Act (7 U.S.C. Sections 2131 - 2159) and the Guide for the Care and Use of Laboratory Animals (8th Edition) as mandated by the U.S. Public Health Service Policy.

All macaques used in this study were housed at the University of Pittsburgh in rooms with autonomously controlled temperature, humidity, and lighting. Animals were singly housed in caging at least 2 square meters apart that allowed visual and tactile contact with neighboring conspecifics. The macaques were fed twice daily with biscuits formulated for nonhuman primates, supplemented at least 4 days/week with large pieces of fresh fruits or vegetables. Animals had access to water ad libitum. Because our macaques were singly housed due to the infectious nature of these studies, an enhanced enrichment plan was designed and overseen by our nonhuman primate enrichment specialist. This plan has three components. First, species-specific behaviors are encouraged. All animals have access to toys and other manipulata, some of which will be filled with food treats (e.g., frozen fruit, peanut butter, etc.). These are rotated on a regular basis. Puzzle feeders, foraging boards, and cardboard tubes containing small food items also are placed in the cage to stimulate foraging behaviors. Adjustable mirrors accessible to the animals stimulate interaction between animals. Second, routine interaction between humans and macaques are encouraged. These interactions occur daily and consist mainly of small food objects offered as enrichment and adhere to established safety protocols. Animal caretakers are encouraged to interact with the animals (by talking or with facial expressions) while performing tasks in the housing area. Routine procedures (e.g., feeding, cage cleaning, etc.) are done on a strict schedule to allow the animals to acclimate to a routine daily schedule. Third, all macaques are provided with a variety of visual and auditory stimulation. Housing areas contain either radios or TV/video equipment that play cartoons or other formats designed for children for at least 3 hours each day. The videos and radios are rotated between animal rooms so that the same enrichment is not played repetitively for the same group of animals.

All animals are checked at least twice daily to assess appetite, attitude, activity level, hydration status, etc. Following M. tuberculosis infection, the animals are monitored closely for evidence of disease (e.g., anorexia, weight loss, tachypnea, dyspnea, coughing). Physical exams, including weights, are performed on a regular basis. Animals are sedated prior to all veterinary procedures (e.g., blood draws, etc.) using ketamine or other approved drugs. Regular PET/CT imaging is conducted on most of our macaques following infection and has proved very useful for monitoring disease progression. Our veterinary technicians monitor animals especially closely for any signs of pain or distress. If any are noted, appropriate supportive care (e.g., dietary supplementation, rehydration) and clinical treatments (analgesics) are given. Any animal considered to have advanced disease or intractable pain or distress from any cause is sedated with ketamine and then humanely euthanized using sodium pentobarbital.

### Research animals

Four cynomolgus macaques (*Macaca fascicularis*), >4 years of age, (Valley Biosystems, Sacramento, CA) were housed within a Biosafety Level 3 (BSL-3) primate facility as previously described and as above. Animals were infected with low dose (∼10 colony-forming units (CFUs)) *M. tuberculosis* (Erdman strain) via bronchoscopic instillation. Infection was confirmed by PET-CT scan at 4 weeks and monitored with clinical and radiographic examinations until 10 weeks post infection.

### Necropsy

Necropsy was performed as previously described^3^. Briefly, an 18F-FDG PET-CT scan was performed on every animal 1-3 days prior to necropsy to measure disease progression and identify individual granulomas. At necropsy, monkeys were maximally bled and humanely sacrificed using pentobarbital and phenytoin (Euthanasia; Schering-Plough, Kenilworth, NJ). Individual granulomas previously identified by PET-CT and those that were not seen on imaging from lung and mediastinal lymph nodes were excised for histological analysis, bacterial burden, and other immunological studies. TB specific gross pathologic lesions and overall gross pathologic disease burden was quantified using a previously published method^102^. The size of each granuloma was measured by pre-necropsy scans and at necropsy. Granulomas were enzymatically dissociated using the gentleMACS dissociator system (Miltenyi Biotec, Inc.) to obtain single cell suspension and used to enumerate bacterial burden and applied on a Seq-Well device for scRNA-seq. 200 μL of each granuloma homogenate were plated in serial dilutions onto 7H11 medium, and the CFU of *M. tuberculosis* growth were enumerated 21 days later to determine the number of bacilli in each granuloma^103^. As a quantitative measure of overall bacterial burden, a CFU score was derived from the summation of the log-transformed CFU/gram of each sample at the time of necropsy.

### Non-human primate single-cell RNA-sequencing (scRNA-seq)

High-throughput scRNA-seq was performed using the Seq-Well platform as previously described^104^. Briefly, total cell counts from single-cell suspension of granuloma homogenate were enumerated and ∼15,000-30,000 cells were applied to the surface of a Seq-Well device loaded with capture beads in the BSL-3 facility at University of Pittsburgh. Following cell loading, Seq-Well devices were reversibly sealed with a polycarbonate membrane and incubated at 37°C for 30 minutes. After membrane sealing, Seq-Well devices were submerged in lysis buffer (5M guanidine thiocyanate, 10 mM EDTA, 0.1% -mercaptoethanol, 0.1% Sarkosyl) and rocked for 30 minutes. Following cell lysis, arrays were rocked for 40 minutes in 2 M NaCl to promote hybridization of mRNA to bead-bound capture oligos. Beads were removed from arrays by centrifugation and reverse transcription was performed at 52°C for 2 hours. Following reverse transcription, arrays were washed with TE-SDS (TE Buffer + 0.1% SDS) and twice with TE-Tween (TE Buffer + 0.01% Tween20). Following ExoI digestion, PCR amplification was performed to generate whole-transcriptome amplification (WTA) libraries. Specifically, a total of 2,000 beads were amplified in each PCR reaction using 16 cycles. Following PCR amplification, SPRI purification was performed at 0.6x and 0.8x volumetric ratios and eluted samples were quantified using a Qubit. Sequencing libraries were prepared by tagmentation of 800 pg of cDNA input using Illumina Nextera XT reagents. Tagmented libraries were purified using 0.6x and 0.8x volumetric SPRI ratios and final library concentrations were determined using a Qubit. Library size distributions were established using an Agilent TapeStation with D1000 High Sensitivity ScreenTapes (Agilent, Inc., USA).

### Non-human primate sequencing and alignment

Libraries for each sample were sequenced on a NextSeq550 75 Cycle High Output sequencing kit (Illumina Inc., Sunnyvale, CA, USA). For each library, 20 bases were sequenced in read 1, which contains information for cell barcode (12 bp) and unique molecular identifier (UMI, 8bp), while 50 bases were obtained for each read 2 sequence. Cell barcode and UMI tagging of transcript reads was performed using DropSeqTools v1.12. Barcode and UMI-tagged sequencing reads were aligned to the *Macaca fascicularis* v5 genome (https://useast.ensembl.org/Macaca_fascicularis/Info/Index) using the STAR aligner. Aligned reads were then collapsed by barcode and UMI sequences to generate digital gene expression matrices with 10,000 barcodes for each array.

### Immunofluorescence staining of macaque granulomas

Formalin-fixed paraffin-embedded granulomas were cut into 5-um sections and deparaffinized and processed as previously indicated using pressure-cooker mediated antigen retrieval and immunofluorescence staining^105,106^. For experiments investigating macrophage protein expression (Fig. S4), we used a cyclic staining process where the antibodies were stripped off the tissue between rounds by running the slide through a cycle of pressure cooking in tris-EDTA buffer as previously described^105,106^. At the end of the multi-round staining process, the tissue section was stripped of antibodies one final time and then stained with hematoxylin and eosin to image the granuloma’s morphologic characteristics. Primary antibodies included CD11b (rabbit polyclonal, Novus Biologicals, Centennial, CO), CD11c (mouse IgG2a, clone 5D11; Leica Microsystems, Deer Park, IL), CD68 (mouse IgG1, clone KP-1; Thermo Fisher Scientific, Waltham, MA), CD163 (mouse IgG1, clone 1D6; Thermo Fisher Scientific), CD206 (mouse IgG2b, clone 685645, Novus Biologicals), FOLR2 (rabbit polyclonal; Novus Biologicals), CSF1R (mouse IgG2b, clone 6B9B9; Novus Biologicals), phospho-SMAD3 (rabbit polyclonal; Novus Biologicals), and phospho-STAT1 (rabbit monoclonal, clone 58D6; Cell Signaling Technology, Danvers, MA). Donkey-anti rabbit or mouse secondary antibodies were purchased from Jackson ImmunoResearch (West Grove, PA). Where possible, multiplexed staining was performed with anti-isotype antibodies purchased from Jackson ImmunoResearch. For pSMAD3 and pSTAT1 staining, CD11c and pSMAD3 were included in a primary antibody cocktail, followed by secondary staining, and then a Zenon rabbit IgG labeling kit (Thermo Fisher Scientific) was used to label the pSTAT1 antibodies to enable the use of two rabbit antibodies in one round of staining. Coverslips were applied with Prolong Gold Mounting medium containing DAPI and the sections were imaged at 20x with a DS-Qi2 camera (Nikon, Melville, NY) on an e1000 epifluorescence microscope (Nikon) operated with Nikon AR Imaging software and acquired as ND2-format images that were exported as TIFF files. For images where multiple rounds of staining were performed, the images were aligned in Adobe Photoshop (Adobe Systems, Mountainview, CA) using the DAPI-stained nuclei for each round as consistent fiducial markers across rounds of staining. For plotting the position of CD11c+ cells expressing combinations pSMAD3 or pSTAT1, the images were segmented with QuPath and data were exported as CSV files for import into CytoMap for analysis and visualization^107,108^. Color schemes were selected to ensure accessibility to all audiences.

### *Ex vivo* macrophage isolation, differentiation, and stimulation

Deidentified buffy coats from three healthy human donors were obtained from MGH Blood Center. PBMCs were isolated from buffy coats by density-based centrifugation using Ficoll (GE Healthcare). Monocytes were isolated using a CD14 positive selection enrichment kit (Stemcell) and frozen in liquid nitrogen. Isolated monocytes were cultured under 10 cytokine conditions, GM-CSF with one of the following cytokines: IFN-β, IFN-γ, IL-1β, IL-4, IL-6, M-CSF, TGF-β, and TNF-α. All cytokines were cultured at 10 ng/mL except for GM-CSF (25 ng/mL) and IL-1β (50 ng/mL). Macrophages were cultured for 1 day, 3 days, or 7 days. Additionally, a separate set of monocytes were differentiated to GM-CSF-derived macrophages then stimulated with the same combination of cytokines on day 6 for 24 hours ahead of RNA-sequencing on day 7. Lastly, on day 3, another set of differentiating macrophages were stimulated with Pam3CSK4 (10 ng/mL). All culture conditions were in RPMI 1640 (ThermoFisher Scientific) supplemented with 10% heat inactivated FBS (ThermoFisher Scientific), 1% HEPES, and 1% L-glutamine.

### *Ex vivo* macrophage RNA-sequencing

RNA-sequencing was performed using prime-seq as described^62^. In brief, cells were lysed in 200 uL of RLT + 1% BME buffer and snap frozen on dry ice. RNA was extracted after proteinase K (15 minutes, 50C) and DNase I digestion (10 minutes, 25C) using SPRI beads. Reverse transcription (RT) was performed by resuspending beads in RT mix and barcoded oligo(dT) primers and incubating 90 minutes at 42C. All samples were then pooled (48 samples per pool) for SPRI-based clean-up, exonuclease digestion, and cDNA amplification. After cDNA amplification, samples were ligated and amplified for sequencing. Libraries were analyzed using Qubit dsDNA HS and Agilent TapeStation D1000 kits. Libraries were sequenced on a NextSeq 500 system (Illumina). Count matrices were generated using kallisto bustools against GRCh38^109^.

### Processing of public datasets

Raw count matrices and metadata from Esaulova *et al.* was accessed via GSE149758^49^. A preprocessed and annotated R object from Pisu *et al.* was downloaded from GSE167232^71^. Skin leprosy and lepromatous lesions/reversal reaction (LL/RR) samples were compiled from GSE150672 and GSE151528, respectively^37,74^. Sarcoidosis samples were compiled from GSE135893^76^. Only sarcoidosis samples were utilized. These single-cell data were processed as described previously. Human lung datasets used for integration are detailed in Table S9.

### Ortholog mapping

Ortholog mapping between human, mouse, cynomolgus macaque, and macaque genomes was performed using the Ensembl database. In analyses where cross-species comparison was utilized, only one-to-one orthologs or genes with identical symbols were included based on the Ensembl database (Ensembl genes 104, Human genes GRCh38.p13) with the following attributes: Gene stable ID, Gene name, Mouse gene name, Mouse gene stable ID, Macaque gene name, Macaque gene stable ID, Crab-eating macaque gene name, Crab-eating macaque gene stable ID.

### Identification of ambient RNA-associated genes

We used SoupX as described to identify potentially problematic genes due to ambient RNA contamination. Ambient contamination per array was automatically estimated (autoEstCont) using the raw count matrix. Gene counts in barcodes not identified as bona fide cells were utilized to determine a list of genes defined as “soup-defining.” The top 100 expressed genes (based on inspection of the distribution of counts within selected arrays) from each array were collated and genes present in at least three arrays with expression levels above the 33rd percentile or genes present in more than 14 arrays were classified as soup-defining. These genes were not included in any PCA or integration analyses. These genes included common housekeeping genes like ACTB, ATP6, COX1, ND6, TMSB4X along with ribosomal genes and dominantly-expressed cell lineage genes.

### Data preprocessing and quality control

Data from cohort 1 (C1) was provided by the authors and is available on the Single Cell Portal (SCP257, SCP1749). From these week 10 data, we extracted originally assigned phagocytes (pDC, cDC, Macrophage, and Mast cell clusters) for downstream analyses. We additionally derived cell type markers from C1 using logistic regression differential expression controlling for the batch covariate for downstream use in annotation. Cohort 2 data were initially filtered through low stringency thresholds (>450 UMIs, >100 genes, <10% mitochondrial reads, <50% ribosomal reads, <10% heat shock family reads, <5 median absolute deviations (MADs)) and clustered. After standard processing (see Data processing, embedding, visualization, and clustering), additional cells were removed based on expert inspection of transcriptional profiles and technical metrics.

### Data processing, embedding, visualization, and clustering

Primary single-cell analyses were performed using Seurat. Counts were log-normalized using NormalizeData and the top 3000 variable features were selected using FindVariableFeatures (selection.method = “vst”). PCA was run on the scaled matrix on variable features only. Selection of downstream PCs was inspected using multiple methods including the “elbow” heuristic and an intrinsic dimension estimation (maxLikGlobalDimEst, intrinsicDimension, R). Batch effects were then corrected using Harmony (theta = 1, sigma = 0.1, lambda = 1, dims.use = 1:30) using 30 PCs. Visualization of the UMAP embedding was generated using RunUMAP across 20 dimensions. Clustering was performed on the shared nearest neighbor (SNN) graph (knn = 20, dimensions = 20) using the Walktrap algorithm (steps = 4, cluster_walktrap, igraph, R) and the Leiden algorithm (leiden_find_partition, leidenbase, R). Any clusters with less than 10 cells were automatically grouped into other clusters based on their SNN connectivity (modified GroupSingletons, Seurat, R). Leiden clustering was performed automatically by scanning resolutions with between 10 and 50 clusters then optimizing the modularity between those resolutions. Leiden and Walktrap results were visually inspected. Resulting clusters were hierarchically clustered and reordered (BuildClusterTree, Seurat, R) based on expression of all variable features. This procedure was standard and utilized across all datasets, as supported by recent benchmarking efforts^110^.

### Cluster annotation

Differentially-expressed genes were calculated using the Wilcoxon Rank Sum Test and AUROC implemented in presto (wilcoxauc) and a logistic regression test implemented in Seurat (FindMarkers) using the array as a latent variable^111^. The log fold-change between the top two expressing clusters was also calculated to more aptly describe gene specificity and expression relative to similar clusters, as described previously^72^. Cells identified as cycling cells were subsetted, reprocessed, and reassigned based on cell type markers. To assist in lymphocyte annotation, we utilized a lung reference as well as original labels for C1^112^. We both scored cells based on differentially-expressed gene signatures and transferred cell labels using *symphony*^113,114^.

### Integration of cohort 1 and cohort 2

Cohort 1 and cohort 2 phagocytes were integrated using the integration procedure in Seurat^111^. Using the reciprocal PCA approach, we integrated all batches with >= 201 cells. Each array was split and processed through PCA (normalization, variable gene identification, scaling, and PCA). Integration anchors were identified using FindIntegrationAnchors (dimensions = 1:30, k.filter = 200, k.score = 20, k.anchor = 5, anchor.features = 3000, n.trees = 20). Data was then integrated using IntegrateData (dimensions = 1:30). Subsequently, integrated data was analyzed as described (see Data processing, embedding, visualization, and clustering). These procedures were performed on a GCP Cloud Compute instance using 64 CPUs and 416 GB.

### Human lung myeloid atlas processing, integration, and analysis

Integration of human lung myeloid cells was performed similarly to NHP cohort integration. All datasets were preprocessed through standardized gene and metadata harmonization. Quality control filters applied include < 20% mitochondrial reads, < 50% ribosomal reads, < 5% hemoglobin or heat shock reads, >= 200 nUMIs and >= 100 genes detected. Mononuclear phagocytes (MNPs) were identified by transferring HLCA labels as described above (see Cluster annotation) and by scoring cells based on signatures derived from that atlas. First, we define a MNP score for each cell, which is the difference between the HLCA scores for MNP cell types and non-MNP cell types. We then fit a Gaussian mixture model to this score and define an upper threshold of 2.5σ above µ. Clusters with a median score above this threshold were labeled as MNPs for the second round of classification. Cycling cells above this threshold were also included.

Clusters that did not reach the median score threshold or contained more than 50% of other cell types (as classified by the maximum Travaglini score) were not MNPs. A second round of preprocessing and annotation revealed contaminating non-MP clusters that were manually inspected and removed. This procedure combining automated labeling procedures and expert curation generated a robust set of mononuclear phagocytes for integration.

We excluded samples with less than 100 cells and pulled the largest 3 control and 3 disease samples from each study. Reference samples were set to the most abundant control and disease sample from each study. Using the reciprocal PCA approach, we integrated all batches with >= 201 cells. Each array was split and processed through PCA (normalization, variable gene identification, scaling, and PCA). Integration anchors were identified using FindIntegrationAnchors (dimensions = 1:20, k.filter = 200, k.score = 20, k.anchor = 5, anchor.features = 2000, n.trees = 20). Data was then integrated using IntegrateData (dimensions = 1:30). These procedures were performed on a GCP Cloud Compute instance using 64 CPUs and 416 GB. Integrated data was then clustered as described previously (see Data processing, embedding, and clustering). CellTypist was used to build a granuloma reference model and predict labels on this atlas^70^. Diversity sampling was performed using scSampler (*96*). Only datasets with both control and disease samples were used. 1,000 cells were sampled across 1,000 iterations for both random and diversity-preserving sampling procedures.

### Comparison with other scRNA-seq datasets

To compare profiles with the mouse scRNA-seq data, we defined gene signatures for each subset in each dataset using methods described above (see Cluster annotation). We subsetted these signatures to 1:1 orthologs and scored subsets using UCell (*AddModuleScore_UCell*). The *Nos2* signature was identified by calculating the Spearman correlation of all genes with *Nos2* and extracting the top 50 genes with a positive Spearman correlation coefficient. To compare with rhesus macaque and human leprosy data, *symphony* was used to build a reference model and transfer labels. Sarcoidosis was compared using the Seurat transfer procedure based on PCA projection across 30 dimensions^111^.

### Enrichment and activity analysis

We utilized Enrichr (enrichR, R package) to perform gene set enrichment analysis on the differentially-expressed genes^115^. The GO Molecular Function database was utilized to calculate enrichment using Fisher’s exact test. We used *decoupleR* to calculate transcription factor and metabolic pathway activity from DoRothEa and KEGG databases, respectively^116^. The normalized weighted mean scoring procedure was used based on its benchmarked performance.

### NicheNet

NicheNet was used to identify potential ligand-receptor activity within myeloid populations, as outlined in the method vignettes^57^. Briefly, we first define sender and receiver populations, background and target gene sets, and potential ligands. Target gene sets were defined as differentially-expressed genes with the highest gene expression across all clusters and an auROC >= 0.6 and adjusted P value <= 0.001. Background gene sets were defined by expression in at least 10% of cells. NicheNet returns ligand activities based on target genes relative to background genes. Potential receptors are then identified from top ligands. In this application, we defined sender populations as cells not utilized for defining the target gene set (e.g., all subsets except RM3). The Pearson correlation coefficient and auROC is reported as a measure of suggested ligand activity.

### Statistical methods

For all the analysis and plots, sample sizes and measures of center and confidence intervals (mean ± SD or SEM), and statistical significance are presented in the figures, figure legends, and in the text. Cellular abundances were tested using a binomial generalized linear model. Inverse Simpson Index was calculated using cell counts (*vegan::diversity* function). Differentially-expressed markers were determined by comparing groups using a Mann-Whitney U test in addition to the auROC metric (wilcoxauc, presto, R). Gene enrichment was calculated using Fisher’s exact test. Score comparisons were conducted using Mann-Whitney U tests, adjusted using the Benjamini-Hochberg procedure. NicheNet statistics were calculated as previously described. All P values and, where appropriate, adjusted P values were considered significant at ≤ 0.05. All statistical analyses were performed in R using base statistics and supporting packages.

## Acknowledgements

We acknowledge the outstanding work of veterinary and research technicians. We thank the Bryson and Blainey lab members for discussions and feedback.

## Author contributions

Conceptualization: JMP, JTM, JLF, AKS, SMF, BDB

Methodology: JMP, HPG, TKH, BFJ, MLN, JTM, JLF, AKS, SMF, BDB

Investigation: JPM, HPG, TKH, BFJ, MLN, JTM, JLF, AKS, SMF, BDB

Data curation: JMP, TKH, CG, SKN, JDB, BFJ, MLN, JTM, JLF, AKS, SMF, BDB, DM

Formal analysis: JMP, CG, BFJ, MLN, JTM, JLF, BDB

Visualization: JMP, BFJ, MLN, JTM, BDB

Resources: DAL, JTM, JLF, AKS, SMF, BDB

Writing - original draft: JMP, PCB, JTM, BDB

Writing - review and editing: all authors Supervision: DAL, PCB, JTM, JLF, AKS, SMF, BDB

Funding: DAL, JTM, JLF, AKS, SMF, BDB

## Data and materials availability

All the data supporting this work is included in the figures, or can be found in the supplementary information, https://fairdomhub.org/studies/1184. Specifically, bulk human MDM RNA-sequencing data have been deposited in the Gene Expression Omnibus (GEO) under accession number GSE211113. Single-cell RNA-sequencing data from non-human primates have been deposited in the Gene Expression Omnibus under accession number GSE211663. For all data analyses, we used publicly available software. All code used for this analysis is available on Github (https://www.github.com/joshpeters).

## Supplemental Figure Captions

**Fig. S1. Technical assessment of ambient and batch effects. (A)** Average expression of genes in non-cellular barcodes across batches colored by ambient thresholds for variable gene exclusion. **(B)** UMAP embedding and LISI metrics for week 10 cohort 2 data colored by batch.

**Fig. S2. Overview of week 10 cohort 2 infection. (A)** UMAP embedding of annotated week 10 cohort 2 data colored by annotated cell type. **(B)** Clustered heatmap of scaled log(TP10K+1) expression values of marker genes across annotated cell types.

**Fig. S3. Inverse Simpson’s Index across sample types. (A)** Inverse Simpson’s Index describing sample diversity across sample types, non-diseased, week 10 early, week 10 late, and week 4. P values were adjusted using the Benjamini-Hochberg procedure. Adjusted P values are denoted by: *, p < 0.05; **, p < 0.01; ***, p < 0.001; ****, p < 0.0001. Only comparisons significant at p < 0.05 are shown.

**Fig. S4. Immunofluorescence staining of myeloid cell markers in macaque granulomas shows that subset-defining antigens are expressed in different granuloma locations.** A cyclic immunofluorescence staining protocol was used on a necrotic granuloma to examine the protein expression and localization of transcriptionally defined subsets identified by scRNA-seq analysis. Protein markers (yellow) are shown against the granuloma’s DAPI-stained nuclei (blue). The inset region represented by the black box shown in the hematoxylin and eosin-stained image (top left) was selected to show representative granuloma regions including intact granuloma-adjacent lung (inset, top left), lymphocyte cuff and epithelioid macrophage regions (inset, middle), and caseum (inset, bottom right).

**Fig. S5. NicheNet analysis of cytokine production.** Cell-type specific analysis of gene expression of cytokines predicted by NicheNet to be acting on RM2 cells.

**Fig. S6. Immunofluorescence staining of pSTAT1 and pSMAD3 in macaque granulomas.** Granulomas harvested from animals euthanized at (A) 4- or (B) 10-weeks post infection were stained for pSTAT1 (green) and pSMAD3 (magenta) as surrogates for IFN-γ and TGF-β signaling, respectively. CD11c (blue) was used as a broadly-expressed macrophage marker (blue) and maps showing the position of the granuloma’s nuclei (grey) and pSTAT1+CD11c+ (green) and pSMAD3+CD11c+ (magenta) macrophages is shown to facilitate visualization of each population’s location (middle panels). The position of three distinct pSTAT1 phenotypes noted on the full granuloma image and zoomed in regions (right) are shown with the region’s nuclei (DAPI; grey) to show the cellularity within each region.

**Fig. S7. Alignment of NHP states with NHP states in Esaulova *et al*. (A)** CellTypist classification of cell populations across this study and Esaulova et al. **(B)** Proportion of predicted granuloma labels from cynomolgus subsets between control, latent, and active rhesus samples.

**Fig. S8. Alignment of NHP states with murine states. (A)** Hierarchically-clustered heatmap of AUROC values for murine subsets annotated from Pisu *et al*. 2020. Supplementary bar plot (*right*) describes the number of DE genes per subset. **(B)** Hierarchically-clustered heatmap of scaled NHP scores of murine signatures. **(C)** Hierarchically-clustered heatmap of scaled murine scores of NHP signatures.

