## Supplementary figures and images for "Systematic deconstruction of myeloid cell signaling in tuberculosis granulomas reveals IFN-γ, TGF-β, and time are associated with conserved myeloid diversity"

### Supplemental Figure 1

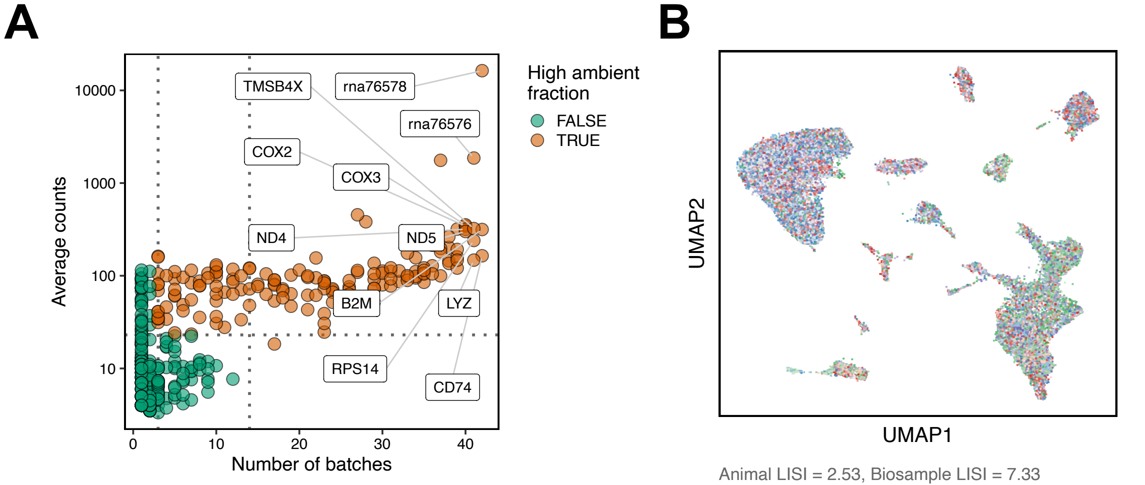

### Supplemental Figure 2

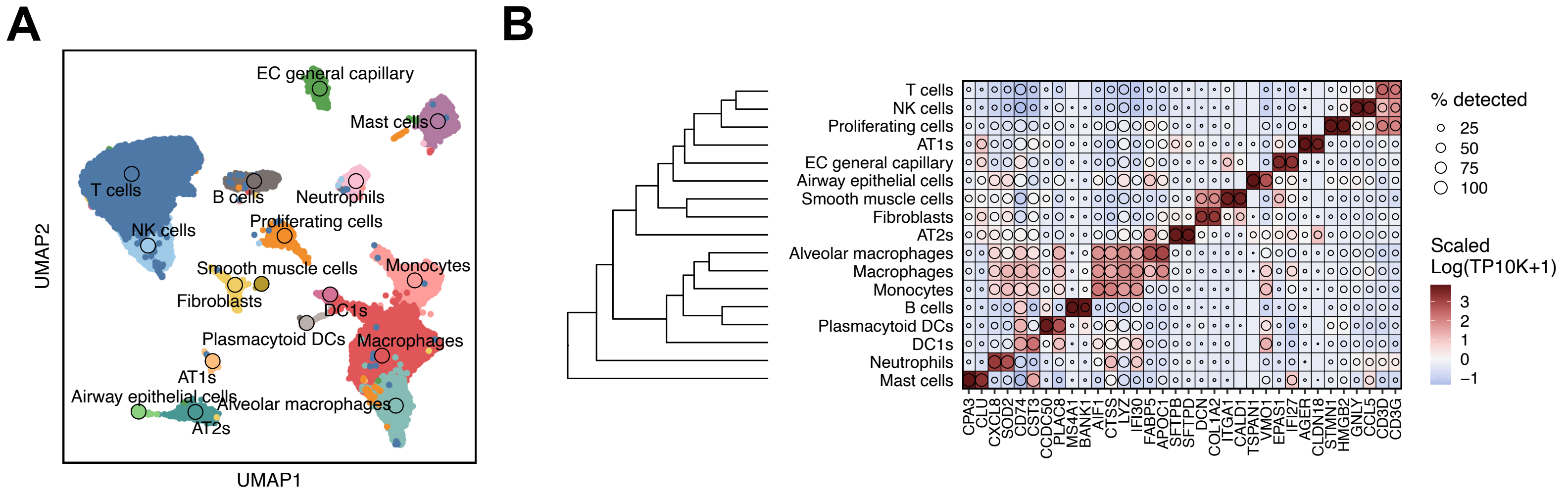

### Supplemental Figure 3

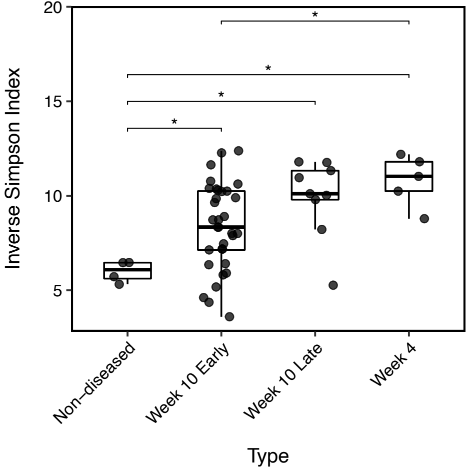

### Supplemental Figure 4

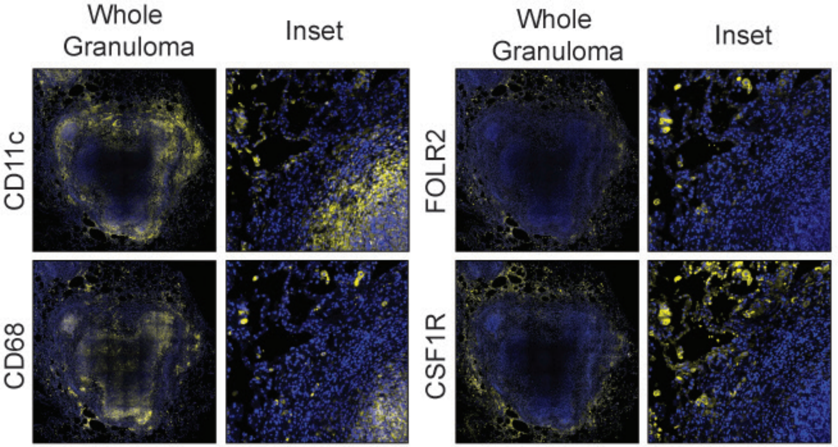

### Supplemental Figure 5

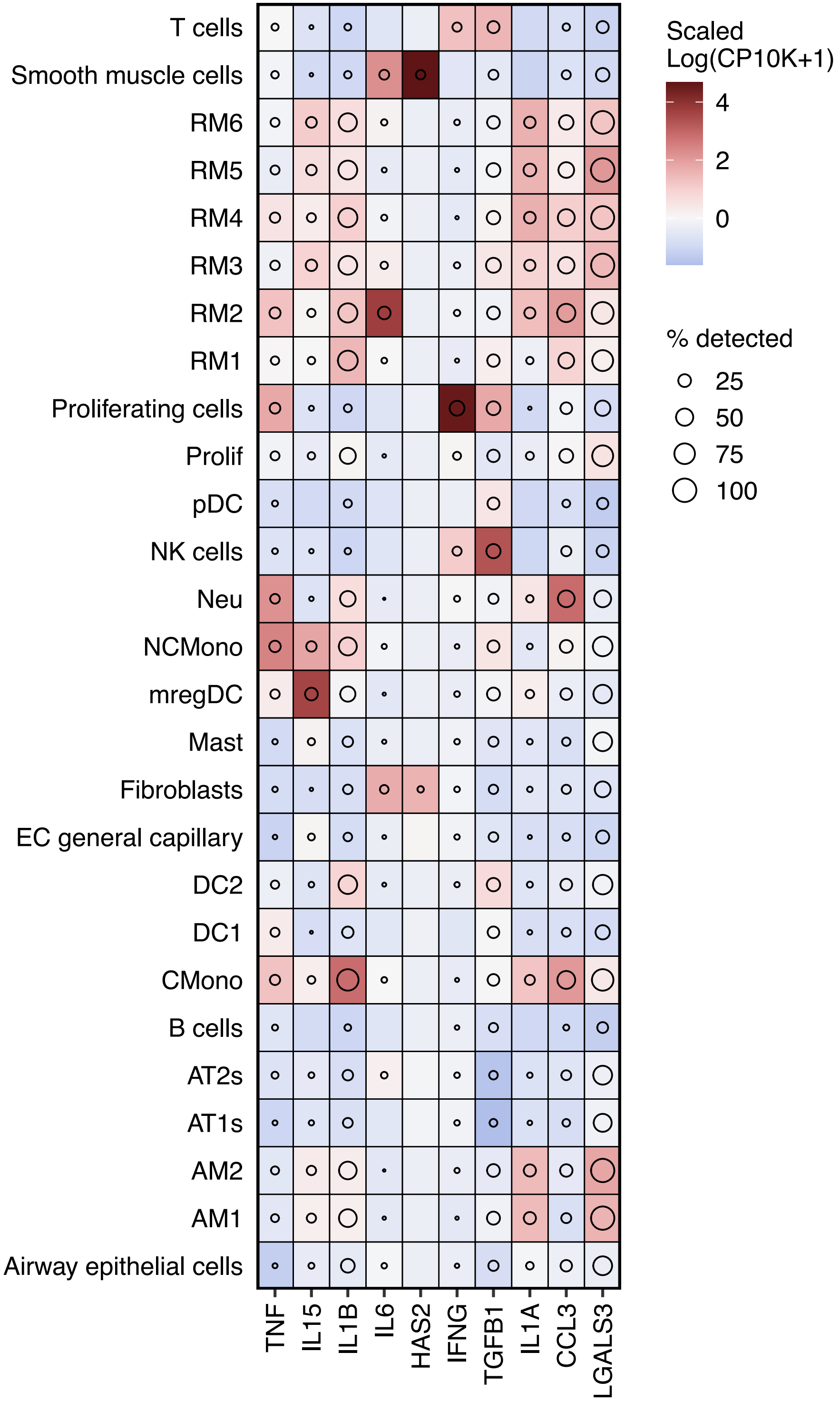

### Supplemental Figure 6

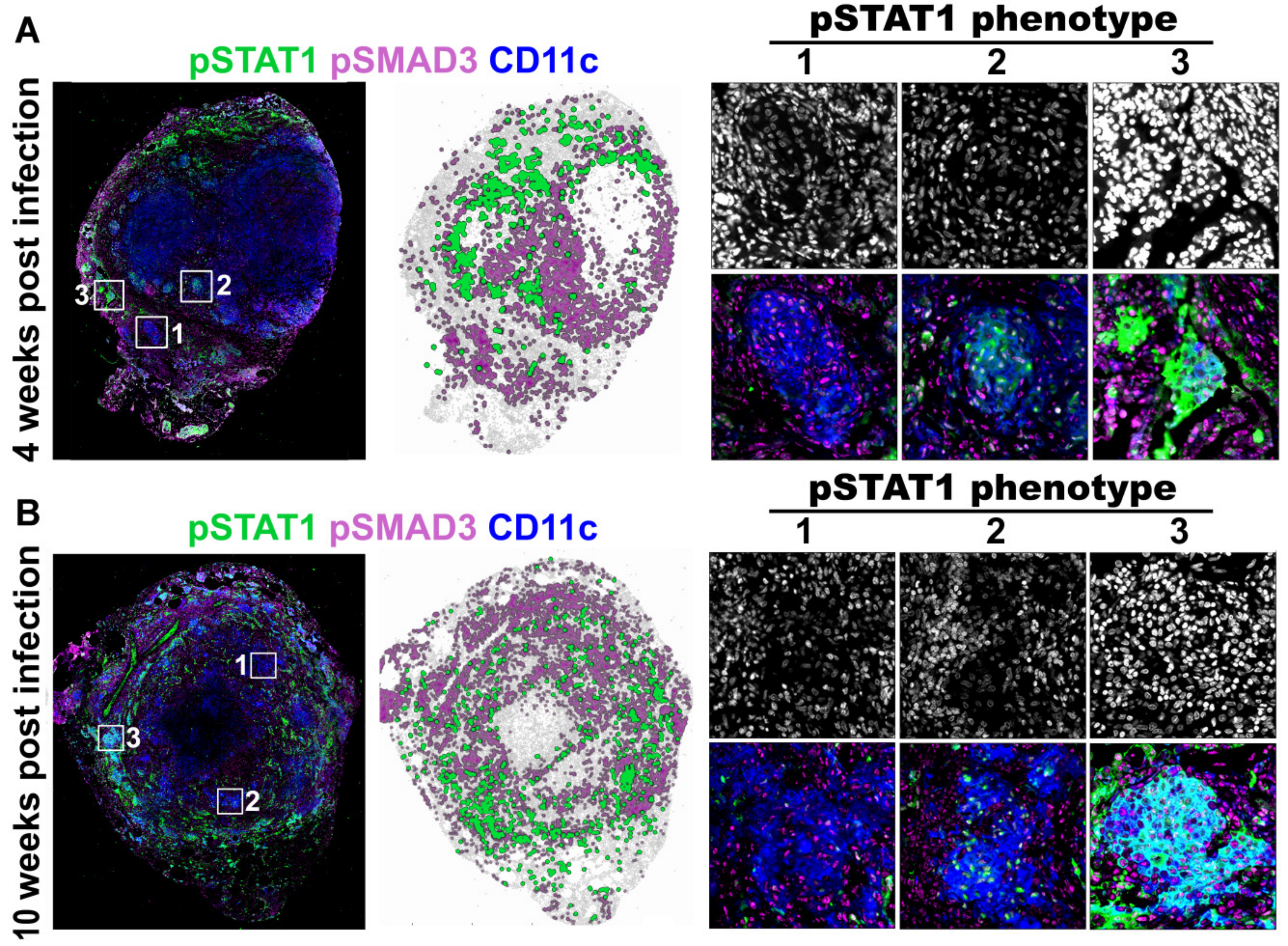

### Supplemental Figure 7

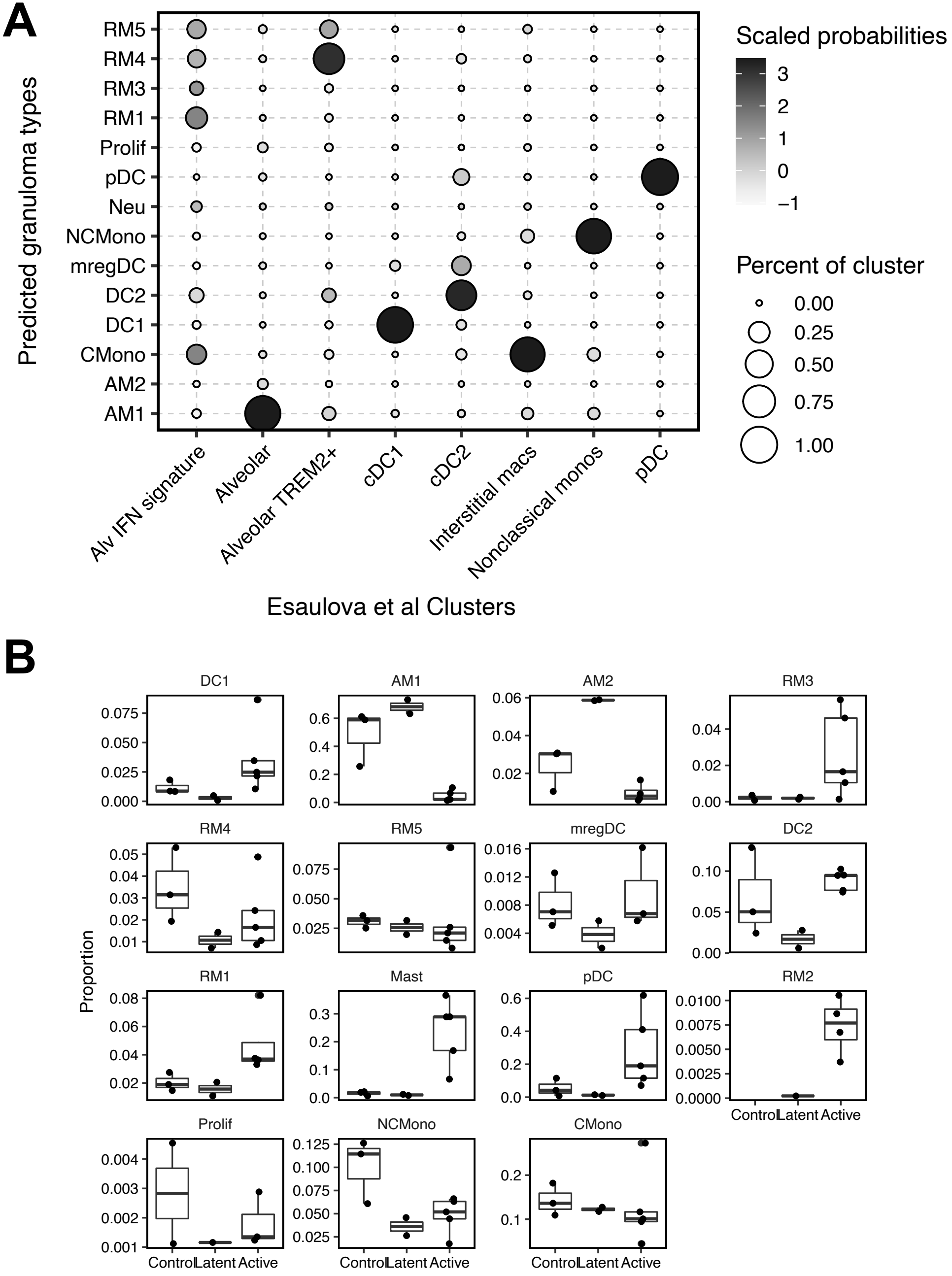

### Supplemental Figure 8

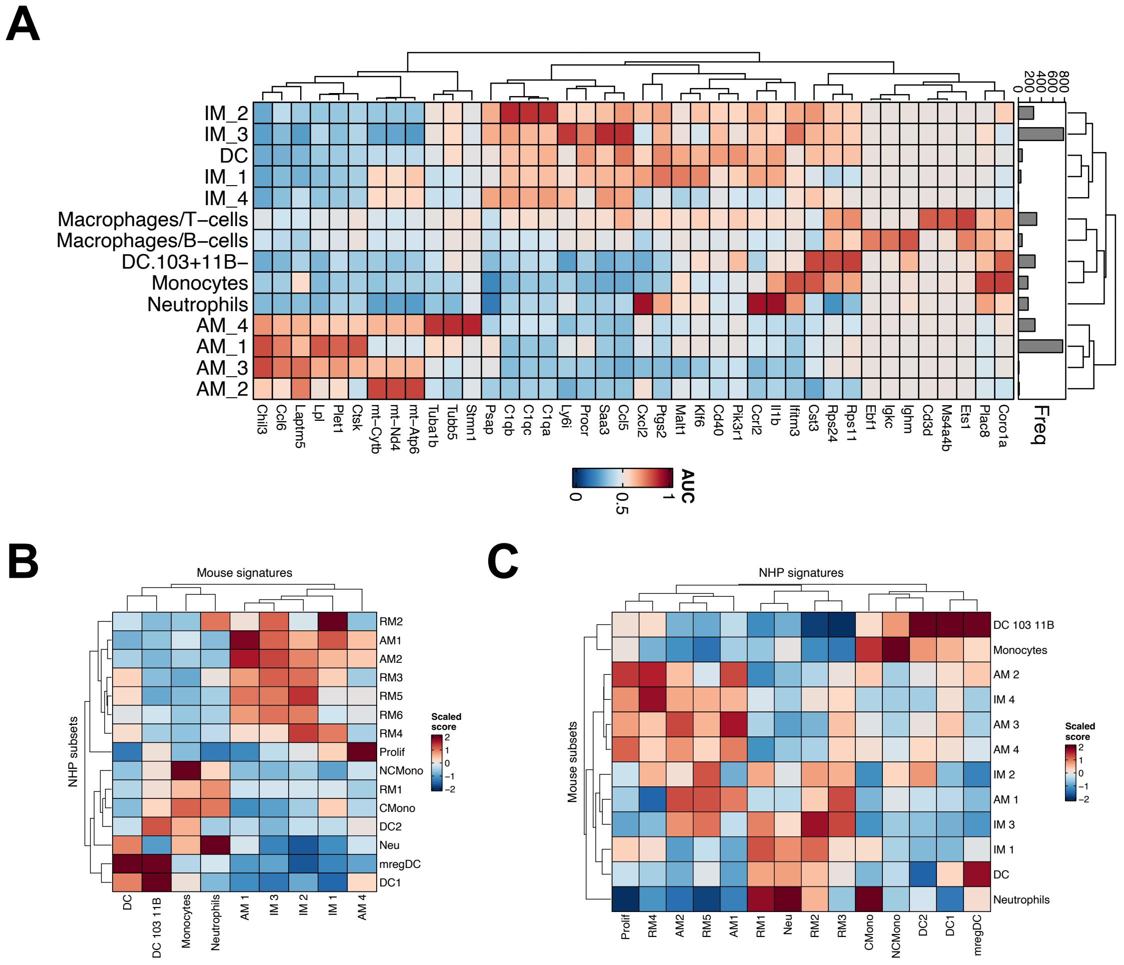
